# Deregulation of LRSAM1 expression impairs the levels of TSG101, UBE2N, VPS28, MDM2 and EGFR

**DOI:** 10.1101/337451

**Authors:** Anna Minaidou, Paschalis Nicolaou, Kyproula Christodoulou

## Abstract

CMT is the most common hereditary neuromuscular disorder of the peripheral nervous system with a prevalence of 1/2500 individuals and it is caused by mutations in more than 80 genes. LRSAM1, a RING finger ubiquitin ligase also known as TSG101-associated ligase (TAL), has been associated with Charcot-Marie-Tooth disease type 2P (CMT2P) and to date eight causative mutations have been identified. Little is currently known on the pathogenetic mechanisms that lead to the disease. We investigated the effect of LRSAM1 deregulation on possible LRSAM1 interacting molecules in cell based models. Possible LRSAM1 interacting molecules were identified using protein-protein interaction databases and literature data. Expression analysis of these molecules was performed in both CMT2P patient and control lymphoblastoid cell lines as well as in *LRSAM1* and *TSG101* downregulated SH-SY5Y cells.*TSG101, UBE2N, VPS28, EGFR* and *MDM2* levels were significantly decreased in the CMT2P patient lymphoblastoid cell line as well as in *LRSAM1* downregulated cells. *TSG101* downregulation had a significant effect only on the expression of *VPS28* and *MDM2* and it did not affect the levels of LRSAM1. This study confirms that LRSAM1 is a regulator of *TSG101* expression. Furthermore, deregulation of LRSAM1 significantly affects the levels of *UBE2N, VPS28, EGFR* and *MDM2*.

## Introduction

Charcot-Marie-Tooth (CMT) disease is the most common inherited neuropathy, characterized by peripheral nervous system degeneration, with a relatively high frequency (1:2,500 individuals)(1, 2). CMT disease is classified into demyelinating, axonal and intermediate types based on electrophysiological findings (3). It is further sub-divided into an increasing number of genetic subtypes with more than 80 CMT genes currently known.

Leucine Rich Repeat And Sterile Alpha Motif Containing 1 (LRSAM1) has been associated with Charcot-Marie-Tooth disease type 2P (CMT2P). LRSAM1, is a RING finger ubiquitin ligase, which participates in a range of functions, including cell adhesion, signaling pathways and cargo sorting through receptor endocytosis (4, 5). It is a multidomain protein with an N-terminal leucine-rich repeat (LRR) domain, followed by an Ezrin-Radixin-Moezin (ERM) domain, a coiled-coil (CC) region, a SAM domain, a PSAP/PTAP tetrapeptide motif and a RING finger domain (6). LRSAM1, also known as TAL (TSG101-associated ligase), regulates the metabolism of *TSG101* (Tumor Susceptibility gene 101) by polyubiquitination (7). *TSG101* is a tumor suppressor gene component of the ESCRT machinery (Endosomal Sorting Complex Required for Transport), with a significant role in cell cycle regulation and differentiation (5, 8). LRSAM1 binds to TSG101, through a bivalent binding of the PTAP motif and the CC domain of LRSAM1 (5, 9). After binding, LRSAM1 attaches several monomeric ubiquitins to the C-Terminal of TSG101, thus regulating its function. LRSAM1 mutations impair the LRSAM1-TSG101 interaction (10).

We identified a dominant *LRSAM1* mutation (c.2047-1G>A, p.Ala683ProfsX3) located in the RING finger domain of LRSAM1 (11). Recently, we reported that downregulation of *LRSAM1* affects the proliferation and morphology of neuroblastoma SH-SY5Y cells, and, overexpression of wild-type *LRSAM1* rescues, while the c.2047-1G>A mutant fails to rescue the phenotype of thecells (12). To date, eight LRSAM1 mutations have been associated with CMT neuropathy, seven of them associated with dominant and one with recessive inheritance (13).

In order to study the role of *LRSAM1* and the effect of the c.2047-1G>A mutant E3 ligase domain, we identified molecules that possibly interact with LRSAM1. Expression levels of selected molecules were evaluated in the *LRSAM1* c.2047-1G>A CMT2P patient lymphoblastoid cell line and also in *LRSAM1* knocked down SH-SY5Y cells. Since, *TSG101* is the only currently well characterized interactor of LRSAM1 (5), we also performed *TSG101* downregulation in SH-SY5Y cells and evaluated the levels of selected molecules in these cells as well.

## Results

### Deregulation of LRSAM1 affects TSG101 expression

Since TSG101 is an already known interactor of LRSAM1, we evaluated TSG101 levels in the CMT2P patient lymphoblastoid cell line. Western blot analysis of the CMT2P patient and control lymphoblastoid cell lines confirmed the presence of the wild-type and the mutant truncated LRSAM1 proteins in the patient sample, as previously reported (4), and revealed a more than 50% reduction of TSG101 levels in the patient lymphoblastoid cell line as compared with the controls (Figure 1).

**Figure 1:**
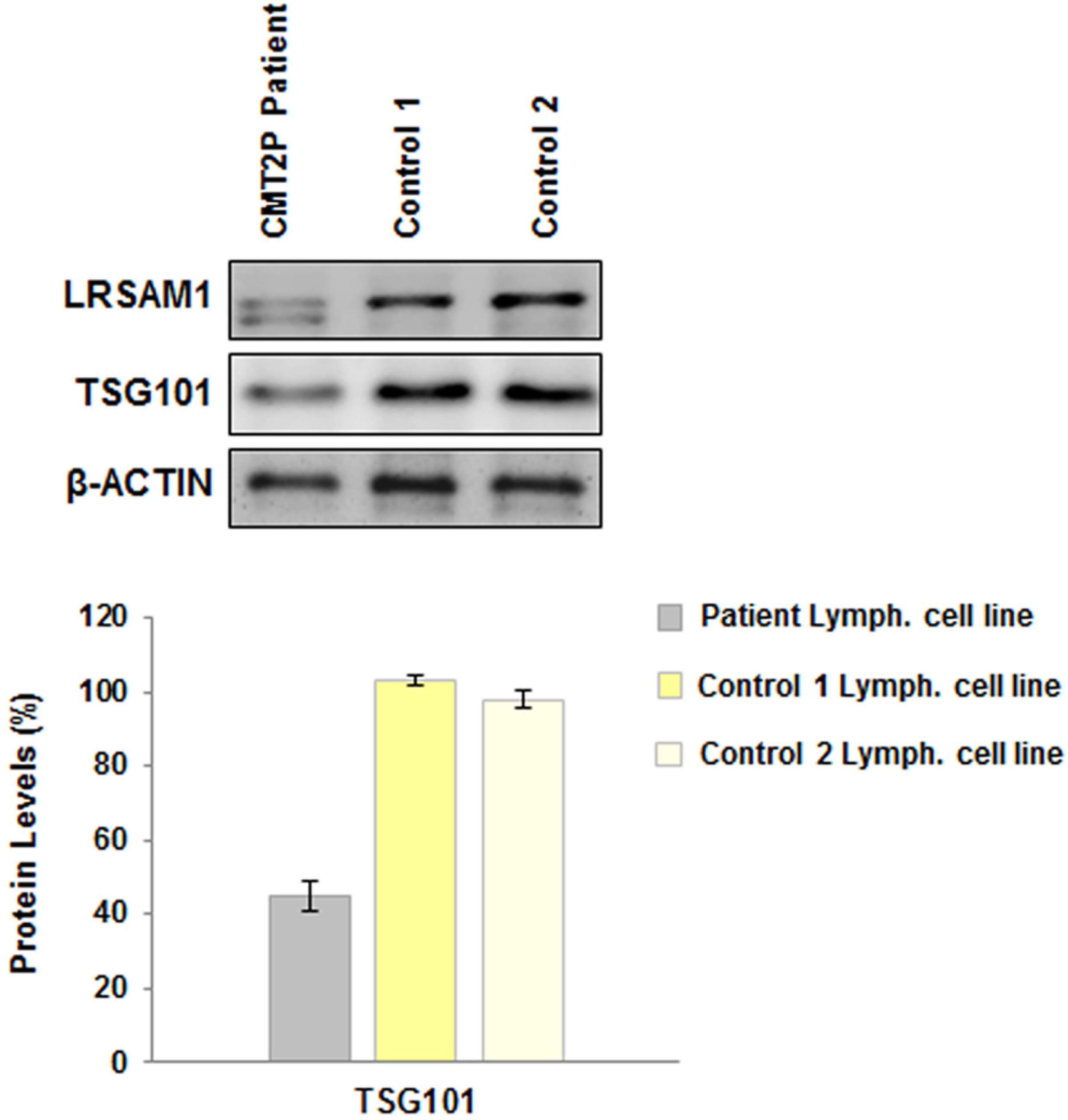
Investigation of TSG101 levels in lymphoblastoid cell lines. Western blot analysis of TSG101 levels in the patient and control (1, 2) cell lines. β-ACTIN was used as an internal control. Quantification of TSG101 was performed relative to β-ACTIN. Control 1 values were set to 100%. **TSG101 levels**: Patient lymphoblastoid cell line (45±4.02%, p<0.03998), Control 1 lymphoblastoid cell line (103±1.24%, p>0.02147), Control 2 lymphoblastoid cellline (98±2.49%).

With another approach, double transfection of *LRSAM1* siRNA in neuroblastoma SH-SY5Y cells caused a >70% decrease of LRSAM1 levels which also resulted to an approximately 50% reduction of TSG101 levels. Double transfection of *TSG101* siRNA caused a 70% decrease on TSG101 levels, while it did not affect the LRSAM1 levels (Figure 2).

**Figure 2:**
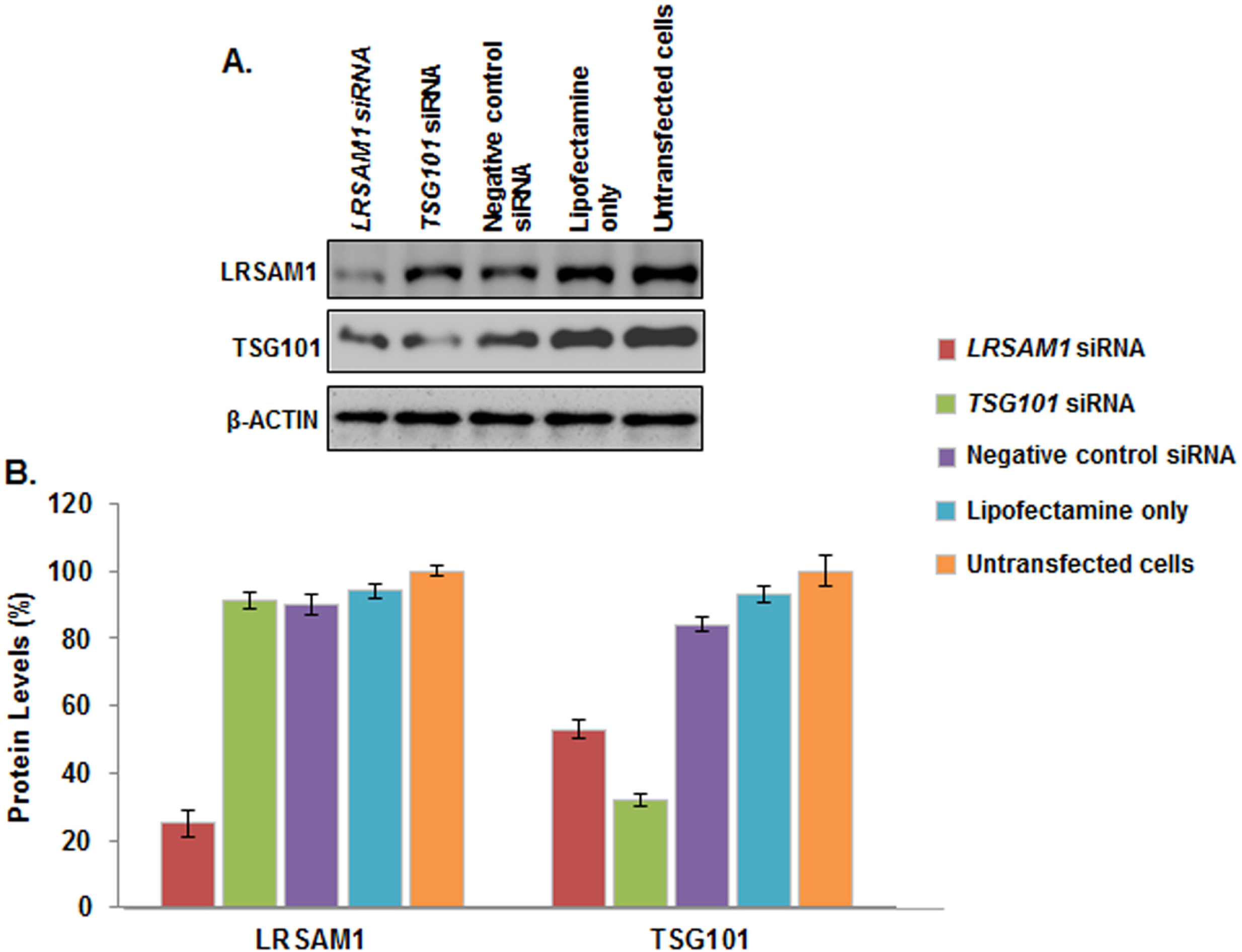
Efficiency of *LRSAM1* and *TSG101* downregulation. (A) Western blot analysis of LRSAM1 and TSG101 levels, compared with the controls (Negative control siRNA, Lipofectamine only, Untransfected cells) 96 hours after the first transfection. β-ACTIN was used as an internal control. (B) Quantification of LRSAM1 and TSG101 was performed relative to β-ACTIN. Untransfected cells values (LRSAM1 and TSG101) were set to 100% and reflect the normal protein levels within the 5 days of the experimental procedure**. LRSAM1 levels**: *LRSAM1* siRNA transfected cells (25±3.94%, p<0.004), *TSG101* siRNA transfected cells (91±2.47%, p<0.0244), Negative control siRNA transfected cells (90±2.89%, p<0.009), Lipofectamine only transfected cells (94±2.15%, p<0.008) and Untransfected cells (100±1.36%). **TSG101 levels**: *LRSAM1* siRNA transfected cells (53±2.72%, p<0.007), *TSG101* siRNA transfected cells (32±1.64%, p<0.0174), Negative control siRNA transfected cells (84±2.20%, p<0.027), Lipofectamine only transfected cells (93±2.49%, p<0.040) and Untransfected cells (100±4.51%).

### Protein-protein interaction network extraction of LRSAM1 possibly interacting molecules

*TSG101, UBE2N, VPS28, EGFR* and *MDM2* that possibly interact with LRSAM1 were identified using the protein-protein interaction database STRING (Figure 3) and IntAct. After literature based evaluation these molecules were selected for further analysis. Furthermore, the AP-3 was selected for analysis since in a recent study LRSAM1 immunoprecipitated with AP-3 (14).

### RNA expression analysis of interacting molecules in lymphoblastoid cell lines

We investigated RNA expression of candidate interacting molecules (*TSG101, UBE2N, VPS28, EGFR, MDM2* and *AP-3*) in the CMT2P patient and control lymphoblastoid cell lines. mRNA levels of *TSG101, UBE2N, VPS28, MDM2* and *EGFR* were significantly reduced in the patient lymphoblastoid cell line as compared to the control, while AP-3 presented an equal expression in both patient and control lymphoblastoid cell lines. Specifically, the patient sample showed significantly lower mRNA levels of *TSG101* (43%), *UBE2N* (45%), *VPS28* (32%), *EGFR* (41%) and *MDM2* (35%) as compared to the control sample. In contrast *AP-3* (98%) showed a similar expression between the patient and control samples (Figure 4).

**Figure 3:**
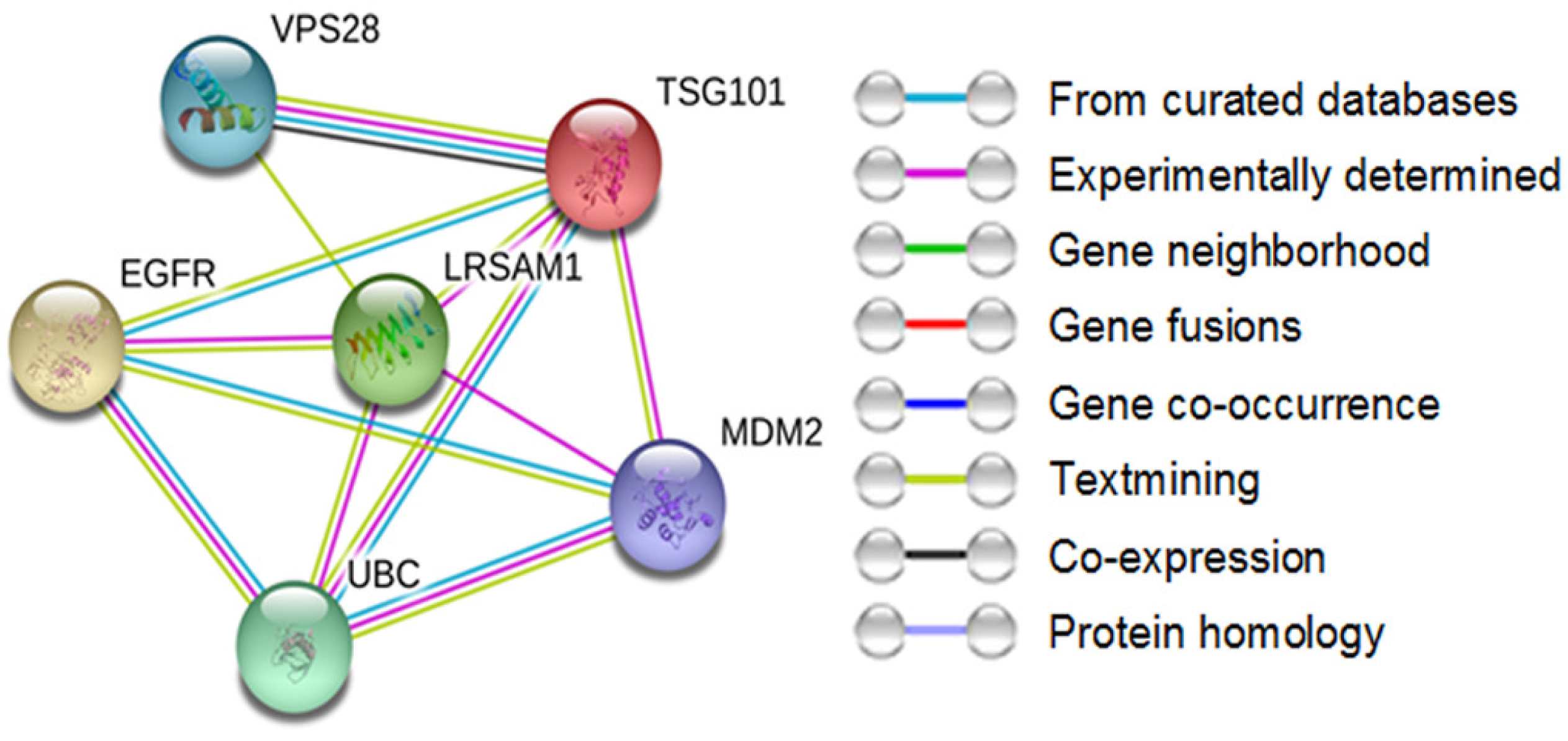
Protein–protein interaction network determined by STRING analysis for LRSAM1 protein. The interaction sources textmining, experiments, databases, co-expression, neighborhood, gene fusion and co-occurrence were used for the selection of the proteins.

**Figure 4:**
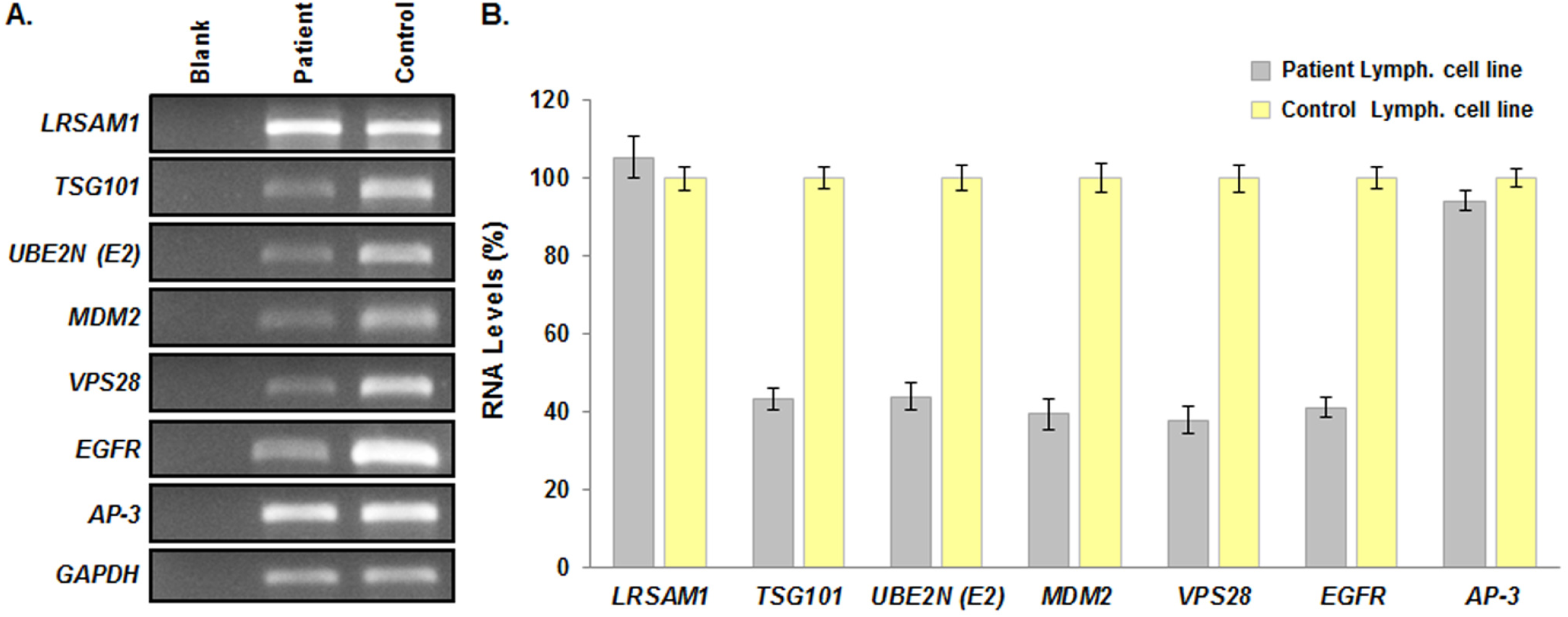
Investigation of LRSAM1 interacting molecules in the CMT2P patient lymphoblastoid cell line. (A) RNA expression analysis of most likely interacting molecules in patient and control samples. *GAPDH* was used as a housekeeping internal control. (B) Quantification of the expression levels of the molecules relative to the control sample, which was set to 100. Values were obtained as ratios of the RNA of interest over *GAPDH* control. Specifically, the patient lymphoblastoid cell line showed significantly lower levels of *TSG101* (43% ±0.28%, p<0.005), *UBE2N* (45%±0.97%, p<0.0109), *VPS28* (32%±3.28%, <0.031), *EGFR* (41%±1.88%, p<0.012) and*MDM2* (35%±2.01%, p<0.025) than the control sample. In contrast *AP-3* (98%±3.47%, p<0.041) showed similar expression between the patient and control. Amplification of each primer set was repeated in triplicate.

### RNA expression analysis of interacting molecules in *LRSAM1* or *TSG101* downregulated SH-SY5Y cells

RNA expression of the above candidate molecules was evaluated in *LRSAM1* or *TSG101* downregulated cells as well. Ninety-six hours after the first *LRSAM1* siRNA transfection, we observed an approximately 70% reduction of endogenous *LRSAM1* RNA levels, reproducing the previously reported effect (5). Downregulation of *LRSAM1* also caused a significant decrease in approximately 50% of *TSG101, UBE2N, VPS28, MDM2* and *EGFR* expression. In agreement with findings in the patient derived lymphoblastoid cell line, *AP-3* levels were not affected by the *LRSAM1* downregultion. Ninety-six hours after the first *TSG101* siRNA transfection, we observed an approximately 70% reduction of endogenous *TSG101* RNA levels. Downregulation of *TSG101* caused an approximately 50% reduction of *VPS28* and a 20% decrease of *MDM2* levels. Normal levels of *LRSAM1, UBE2N, EGFR* and *AP-3* were observed after *TSG101* down regulation (Figure 5). Although, *LRSAM1* downregulation caused a significant reduction of TSG101 levels (55%), *TSG101* downregulation did not affect the expression of *LRSAM1.*

**Figure 5:**
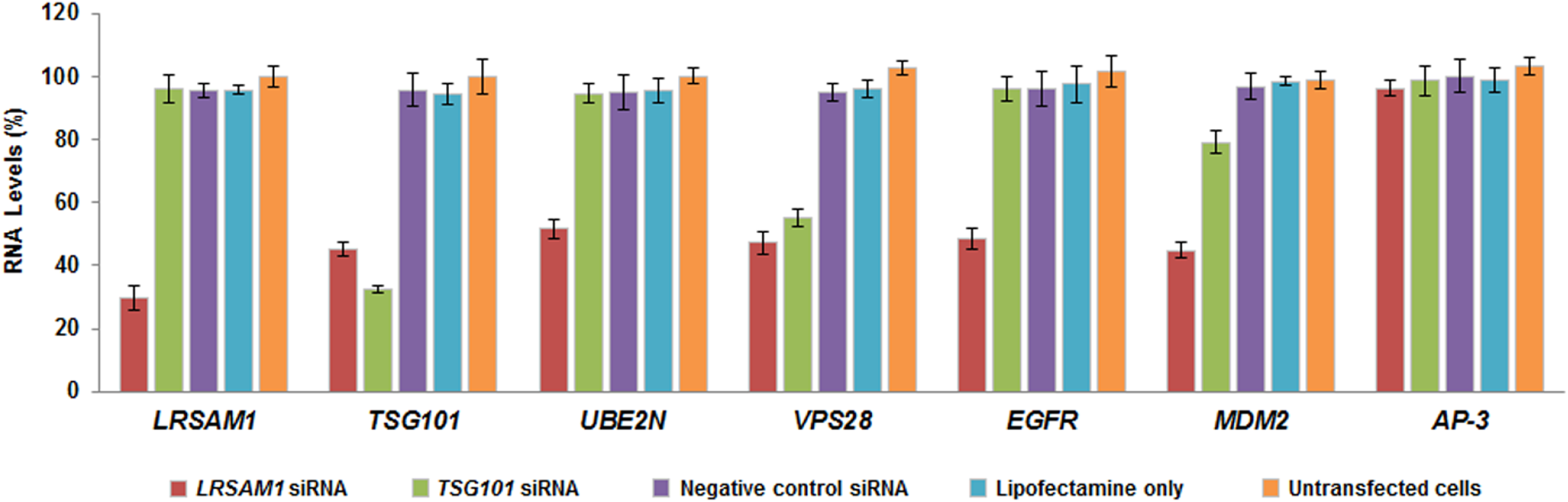
Investigation of LRSAM1 interacting molecules in *LRSAM1* or *TSG101* downregulated cells. Quantification of the RNA expression levels of the interacting molecules relative to the Untransfected control cells, which was set to 100 96h after the first siRNA transfection. Values were obtained as ratios of the RNA of interest over *GAPDH* control. Specifically, ***LRSAM1* downregulation** significantly decreased the levels of *LRSAM1* (30%±3.86%, p<0.0275), *TSG101* (45% ±2.19%, p<0.0092), *UBE2N* (52%±3.05%, p<0.0158), *VPS28* (47%±3.62%, p<0.0336), *EGFR* (49%±3.38%, p<0.0125) and *MDM2* (45%±2.44%, p<0.0096) compared to the Untransfected control cells. In contrast, *AP-3* (96%±2.59%, p<0.0109) showed similar expression between the *LRSAM1* knocked down and Untransfected cells. ***TSG101* downregulation** significantly decreased the levels of *TSG101* (33%±1.11%, p<0.0068) and *VPS28* (55%±2.86%, p<0.0154) and also caused a small decrease of *MDM2* (79%±3.37%, p<0.0217) compared to the Untransfected cells. Expression levels of *LRSAM1* (96% ±4.41%, p<0.0427), *UBE2N* (94%±2.97%, p<0.0172), *EGFR* (96%±3.96%, p<0.0298) and *AP-3* (99%±4.57%, p<0.0462) were similar to the Untransfected cells.

## Discussion

LRSAM1 has been associated with CMT disease and currently its role in neurodegenerative disorders and in CMT pathogenesis is still unclear. In the present study we found that deregulation of *LRSAM1* impaired *TSG101, UBE2N, VPS28, MDM2* and *EGFR* levels. *TSG101* downregulation had an effect on *VPS28* and *MDM2* levels.

The E3 ligase activity of LRSAM1 performs ubiquitination of TSG101, regulating its expression (5). TSG101 levels are also modulated by other mechanisms beyond LRSAM1 E3 regulation. MDM2, another E3 ligase, shares an autoregulatory loop with TSG101 (15). TSG101 prevents its ubiquitination and increases the levels of MDM2 thus promoting ubiquitin degradation of p53 (15, 16). Over expression of truncated TSG101 in U20S cells caused reduction of MDM2 levels (15), while over expression of normal TSG101 doubled the short half-life of MDM2 and elevated its expression levels (15). Notably we detected that 70% downregulation of LRSAM1 caused 55% decrease in TSG101 and also 55% decrease in MDM2, whereas 70% downregulation of TSG101, which did not affect LRSAM1, caused only 20% decrease in MDM2 expression. These results indicate that TSG101 exerts a small effect on MDM2 expression (20%) relative to the effect of LRSAM1 (55%), the later through an unknown mechanism.

Interaction of TSG101 (yeast VPS23) with VPS28, VPS37 and MVB12 is necessary for the formation of the ESCRT-I complex (17). Binding between TSG101 and ESCRT-I subunits protects TSG101 degradation (9). VPS28 binds to the C-Terminal of TSG101 and blocks the interaction with LRSAM1, thus subsequently decreases the ubiquitination of TSG101 (18). Reduced levels of VPS28 were observed in cells that lack TSG101 (18), while over expression of VPS28 did not affect the levels of TSG101 (9). In our study, we observed decreased levels of VPS28 in the CMT2P patient lymphoblastoid cell line and showed that down regulation of LRSAM1 or TSG101 caused a significant reduction in VPS28 levels. It is thus possible that mutant or knockdown LRSAM1 affects VPS28 levels directly or through the impaired TSG101 levels. TSG101 and/or VSP28 deficiency possibly affect the formation of the ESCRT-I complex. The ESCRT-I complex along with ESCRT-0, ESCRT-II and ESCRT-III form the endosomal sorting complexes required for transport (ESCRTs). Each ESCRT complex, consists of VPS proteins, and regulates biogenesis of the Multivesicular Body (MVB) and cargo sorting within the lysosomes. Besides endocytic trafficking, tumor suppression, autophagy, miRNA function and cancer, the ESCRT has been associated with neurodegenerative diseases (19, 20). In the central nervous system endosomal trafficking and lysosomal functions are necessary processes for the survival of neurons. A recent study showed that, lack of Hrs (component of ESCRT-I) caused accumulation of ubiquitinated protein and neurodegeneration in the central nervous system of a mouse model (21). In addition, mutated CHMP2B (component of ESCRT-III) leads to Frontotemporal dementia linked to chromosome 3 (FTD-3) because of autophagosome and ubiquitinated protein accumulation (20). It is thus possible that dysregulation of any ESCRT complex can affect the normal function of the ESCRT machinery and the central nervous system.

We observed a decreased expression of *UBE2N* in the patient lymphoblastoid cell line and also in *LRSAM1* downregulated cells. A recent study showed, using a ubiquitination assay, that LRSAM1 interacts with UBC13, a mouse homolog of UBE2N and that mutations of LRSAM1 prevent this interaction, abolishing the ubiquitination activity of LRSAM1 (10). UBE2N belongs to the family of Ubiquitin-conjugating enzymes (E2) and it is the only known enzyme that facilitates the creation of K63-linked polyubiquitination chains on proteins which are selected for proteasomal degradation (22, 23). E2 along with E1 and E3 enzymes consist the UPS system which removes the misfolded and abnormal proteins within the cells. Alzheimer, Parkinson and Huntington neurodegenerative diseases are caused by the formation of inclusions in the brain because of the accumulation of misfolded proteins in neurons (23), indicating a possible pathogenic role of the UPS system in neurodegeneration. Expression of UBE2N was found to be elevated in synaptosomes in tissues obtained from aged monkey brain while UPS activity was found to be depressed. The increased level of UBE2N in aged monkey brain or the over expressed UBE2N in HEK293 cells, caused the accumulation of mutant huntingtin (HTT). In contrast, the decreased level of UBE2N in aged monkey brain, HEK293 and PC12 cells caused reduction of mutant HTT accumulation (23). Knockdown of UBE2N in Parkin transfected HeLa cells inhibited the K63-linked polyubiquitination at mitochondrial sites suggesting a contribution of UBE2N in Parkin-mediated mitophagy (24).

The RING finger domain of LRSAM1 also plays a role in the regulation of EGFR expression (5). We observed that the LRSAM1 c.2047-1G>A mutation, located in the RING finger domain, results in impaired EGFR expression. Transfection of an inactive mutant-RING LRSAM1 has been reported to lead to the reduction of EGFR expression, while transfection of wild type LRSAM1 improved EGFR expression (5). Also in the presence of mutant TSG101 in NIH 3T3 and SL6 cells, EGFR moved from the early endosomes and MVBs to the plasma membrane for recycling (25). In our study, downregulation of TSG101 did not affect the expression of EGFR. The EGF receptor is a family member of the ErbB receptors of kinases and regulates essential signaling pathways including cell proliferation and differentiation, cell cycle and migration (26). Upregulation of EGFR has been associated with both oncogenesis and neurodegeneration. Expression of EGFR is activated in many malignancies such as breast cancer, prostate cancer and gliomas (27) and stimulates growth and differentiation of neurons (28). Increased levels of the EGFR ligand promoted neuronal death, while high EGFR immunoreactivity was detected in neurites near the neuritic plaques of Alzheimer disease. Lack of Presenilin (PS1), a regulator of EGFR expression, increased EGFR levels resulting to abnormal cell cycle re-entrance (27, 28).

AP-3 has been reported to form a heterotetrameric complex, targeting cargos of lysosomes and lysosome related organelles (29). Recently LRSAM1 was identified as an AP-3 interactor, because it immunoprecipitated with AP-3. It has been proposed, that while LRSAM1 is bound to AP-3, it regulates the ESCRT-I sorting function, by ubiquitinating certain cargoes (14). We found normal expression of AP-3 in the CMT2P patient lymphoblastoid cell line as well as after LRSAM1 downregulation in SH-SY5Y cells. These data, suggest that the interaction between the two molecules is probably controlled by AP-3 and not LRSAM1.

In conclusion, we confirm that *TSG101* expression is regulated by LRSAM1. Decreased levels of *UBE2N* and *EGFR* were observed only after *LRSAM1* knockdown thus indicating that these two molecules are regulated by a mechanism in which only LRSAM1 is involved. Decreased expression of *VPS28* and *MDM2* after either *LRSAM1* or *TSG101* downregulation demonstrates that the levels of these two molecules are regulated by a mechanism in which both LRSAM1 and TSG101 may be involved. Further investigation of the above molecules may enable delineation of the pathways that lead to CMT2P pathology.

## Materials and Methods

In this study, we selected possible LRSAM1 interacting molecules and investigated their expression levels in lymphoblastoid cell lines as well as in *LRSAM1* and *TSG101* downregulated neuroblastoma SH-SY5Y cells.

### Cell culture

#### a. Lymphoblastoid cell cultures

Lymphoblastoid cell lines were established from a CMT2P patient and two normal control individuals using peripheral blood. Lymphocytes were collected from peripheral blood using Ficoll-Paque Plus (Sigma-Aldrich, USA). Selected lymphocytes were infected with the Epstein-Barr virus (EBV) and were cultured in DMEM medium supplemented with 2% FBS.

#### b. Human SH-SY5Y neuroblastoma cells culture

Human SH-SY5Y neuroblastoma cells (ECACC, Sigma-Aldrich, U.S.A), were cultivated in Dulbecco’s Modified Eagle Medium DMEM (Invitrogen, U.S.A.) growth medium without L-glutamine. The DMEM medium was supplemented with 10% FBS (Invitrogen, U.S.A.), 2% GlutaMAX^^TM^^ (Gibco, U.S.A.) and 1% Penicillin-Streptomycin 100X Solution (Invitrogen, U.S.A.). Medium was changed every 2 or 3 days and 0.25% Trypsin-EDTA (Life Technologies, U.S.A.) was used for routine splitting of the cell culture. Both cell lines were incubated in a humidified atmosphere under 5% CO2 at 37°C.

### Whole human LRSAM1 constructs

The pIRES2-EGFP-*LRSAM1* wild-type and mutant constructs were purchased from Eurofins (Germany) as previously descripted (12). The mutant *LRSAM1* cDNA construct included a G base deletion at the first base of exon 25, creating the frame shift at the RNA level (11).

### Downregulation of *LRSAM1*or *TSG101* in neuroblastoma SH-SY5Y cells

Transfections were performed using the Lipofectamine^®^ 3000 (C3019H, Life Technologies, U.S.A.) The siRNAs against *LRSAM1* or *TSG101* (Life Technologies, USA) were double-transfected into SH-SY5Y cells as previously descripted (12) and according to the manufacturer’s instructions. The appropriate amount of siRNA and Lipofectamine^®^ 3000 were dissolved separately in the Opti-MEM^®^ reduced serum medium (Life Technologies, U.S.A.) without FBS and antibiotics. Negative control siRNA (Life Technologies, USA), lipofectamine only and untransfected cells were used as controls of the experiments. Twenty-four hours after each transfection, medium was replaced with fresh DMEM medium. Cells were harvested 96 hours after the first transfection for protein and RNA extraction.

### Protein-protein interaction database

In silico analysis was carried out using the STRING9.05&10.0 (http://string-db.org/) and IntAct (http://www.ebi.ac.uk/intact/) databases. Extracted LRSAM1 possibly interacting molecules were selected for RNA expression analysis after literature evaluation.

### RNA isolation and cDNA synthesis from experimental SH-SY5Y cells and lymphoblastoid cell lines

Cells were collected in PBS and total RNA was isolated using the Qiagen RNeasy kit (Qiagen, U.S.A.). 1% β-Mercaptoethanol (Sigma-Aldrich, U.S.A.) was added in lysis buffer before use. Whole cDNA was synthesized using the ProtoScript^®^ First Strand cDNA Synthesis Kit (New England Biolabs, U.K.) using the oligo-dT primer d (T)23VN according to the manufacturer instructions.

### RNA expression levels

Expression levels of *TSG101, UBE2N, VPS28, MDM2, EGFR* and *AP-3* were evaluated by cDNA PCR amplifications. At least two sets of primers specific for each molecule and also for the *GAPDH* housekeeping gene were used. For each set of primers a standard curve was plotted in order to define the optimum number of PCR reaction cycles that fall within the exponential phase of the amplification reaction (Supplementary Figure 1, available upon request). RNA levels of the patient and control samples as well as of the LRSAM1 and TSG101 knocked down cells were normalized using the *GAPDH* housekeeping RNA levels. Gel bands were measured using the ImageJ analysis software.

### Protein extraction from experimental SH-SY5Y cells and lymphoblastoid cell lines

Proteins were extracted using a protein lysis buffer, that contained 1M NaCl, 10mM Tris-Cl (pH=7.5), 10% glycerol (G5516, Sigma-Aldrich, U.S.A), 1% Tween^TM^20 (Affymetrix, U.S.A.), 10Mm β-Mercaptoethanol (Sigma-Aldrich, U.S.A) and 1x EDTA-free Protease Inhibitor Cocktail (Promega, U.S.A.). Cell lysates were centrifuged at 4°C and supernatants were collected. Lysates were sonicated and denatured at 95°C. Then proteins were diluted in 1x Sodium Dodecyl Sulfate 20% (ThermoFisher Scientific, U.K.). The Coomassie Plus (Bradford) protein assay (23236, ThermoFisher Scientific, U.K.) was used to determine the protein concentration.

### Western blot analysis

Proteins were loaded and separated onto SDS-PAGE 12% polyacrylamide gels and were transferred to Hybond-P hydrophobic polyvinylidene difluoride (PVDF) membranes (Millipore, Germany). Membranes were blocked for 1 hour at room temperature and were incubated overnight at 4°C with the respective specific primary antibody for each protein [mouse anti-LRSAM1/Abcam ab73113 (1:400), rabbit anti-LRSAM1/Novus Biological H00090678-D01 (1:750), mouse anti-TSG101/Novus Biological NB200-11 (1:500) and mouse anti-β-ACTIN/Sigma-Aldrich A2228 (1:4000)]. Then, incubation with the appropriate secondary antibodies was performed for 2 hours and followed by incubation with the visualization LumiSensor^^TM^^ Chemiluminescent HRP Substrate Kit (Genscript, U.S.A.) at room temperature. Membranes were visualized using the UVP imaging system (BioRad, U.S.A.). Quantification of Western blots was performed using the ImageJ program (https://imagej.nih.gov). The protein quantity ratio was estimated relative to β-Actin.

### Statistical analysis

Quantitative data (ratio %) from three independent experiments were analyzed using the two-tailed Student’s paired t-test. Thresholds of P-value < 0.05 were considered statistically significant. All the data are expressed as the mean% ± standard deviation (SD) from three independent experiments. The mean of quantitative data of the control samples was set to 100%.

## Acknowledgements

This work was supported by a grant of the Cyprus TELETHON (Grant number: 73115).

**Abbreviations**
AP-3: Adaptor Protein 3
CC: Coiled-Coil
CMT: Charcot-Marie-Tooth disease
CMT2: Axonal type2
DMEM: Dulbecco’s Modified Eagle Medium
FBS: Fetal Bovine Serum, FBS
E1: Ubiquitin-activating enzyme
E2: Ubiquitin-conjugating enzyme
E2-Ub: Ubiquitin-conjugating enzyme, ubiquitin
E3: Ubiquitin protein ligase
EBV: Epstein-Barr Virus
EGFP: Enhanced green fluorescent protein
EGFR: Epidermal Growth Factor Receptor
ERM: Ezrin-Radixin-Moezin
ESCRT-I: Endosomal Sorting Complex Required for Transport I
HTT: Huntingtin
LRR: N-terminal leucine-rich repeat
LRSAM1: Leucine Rich Repeat and Sterile Alpha Motif Containing 1
MAPKs: Mitogen activated protein kinase signaling
MDM2: Mouse Double Minute 2 homolog
MVB: Multivesicular Body
MVB12: Multivesicular Body protein 12
NF-κB: Kappa B pathway
S-box: C-terminal Steadiness box
TAL: TSG101-Associated Ligase
TSG101: Tumor Susceptibility Gene 101
UBE2N: Ubiquitin-conjugating Enzyme E2N
VPS28: Vacuolar Protein Sorting 28

